# Repeatable ecological dynamics govern response of experimental community to antibiotic pulse perturbation

**DOI:** 10.1101/2020.03.10.985184

**Authors:** Johannes Cairns, Roosa Jokela, Lutz Becks, Ville Mustonen, Teppo Hiltunen

## Abstract

In an era of pervasive anthropogenic ecological disturbances, there is a pressing need to understand the factors constituting community response and resilience. A detailed understanding of disturbance response needs to go beyond associations and incorporate features of disturbances, species traits, rapid evolution and dispersal. Multispecies microbial communities experiencing antibiotic perturbation represent a key system with important medical dimensions. However, previous microbiome studies on the theme have relied on high-throughput sequencing data from uncultured species without the ability to explicitly account for the role of species traits and immigration. Here we serially passaged a 34-species defined bacterial community through different levels of pulse antibiotic disturbance, manipulating the presence or absence of species immigration. To understand the ecological community response measured by amplicon sequencing, we combined initial trait data measured for each species separately and metagenome sequencing data revealing adaptive mutations during the experiment. We found that the ecological community response was highly repeatable within the experimental treatments, owing to an increasingly strong yet canalized response at increasing antibiotic levels, which could be partly attributed to key species traits (antibiotic susceptibility and growth rate). Increasing antibiotic levels were also coupled with increasing species extinction probability, making species immigration preventing this critical for community resilience. Moreover, we could detect signals of antibiotic resistance evolution occurring within species at the same time scale, leaving evolutionary changes in communities despite recovery at the species compositional level. Together these observations reveal a disturbance response which appears as classic species sorting but is nevertheless accompanied by rapid within-species evolution.

## Introduction

In the Anthropocene^1^ characterized by anthropogenic perturbations of environments ranging in scale from the individual organism (e.g. gut microbiota of mammal) to the global ecosystem (e.g. climate change and loss of biodiversity), it is paramount to understand factors contributing to biological resilience^2^. Understanding how the collateral effects of perturbations percolate through the ecosystem, with possibly unintended consequences, is vital to better understand the risks and benefits of human driven control efforts in the restoration and conservation of populations. For instance, antibiotic treatment affects not only the pathogen population but also off-target species in the microbiota of the patient, promoting the spread of antimicrobial resistance^3^. Rational interventions require a detailed, ideally mechanistic, understanding that goes beyond associations, integrating community dynamics, species traits, environmental variables, evolutionary events and stochasticity. However, we are far from such an understanding, which has in part been attributed to the sparsity of controlled studies amidst *in vivo*, field and theoretical studies^4^. Nevertheless, recent advances in predictive modeling suggest that such an understanding is possible for certain rapidly evolving systems^5^.

A notable case of interest is the response of multispecies bacterial communities to perturbations by antibiotics, pharmaceuticals and other compounds of an anthropogenic origin. Understanding this response can be critical, among others, for rational therapeutics to mitigate unwanted effects on patient health caused by changes in the gut microbiota (e.g. *Clostridioides difficile* infection^6^), management of waste water treatment to ensure the maintenance of key functionalities in bacterially driven processes^7^, and redesigning of agricultural practices to maintain microbiota contributing to crop health and productivity^8^.

Traditionally, ecological timescales have been considered shorter than evolutionary timescales, causing ecological processes to drive the community response to environmental change. The community response to perturbation is therefore expected to be determined, similar to other forms of community assembly, by environmental filtering on pre-existing traits, particularly at the species level^9^. The disturbance response of multispecies microbial communities may also be likened to the selective response of heterogeneous populations of single species, where standing variation leads to repeatability and predictability^10-13^. However, the role of standing trait variation at the species level in the bacterial disturbance response remains unclear, since large population sizes and short generation times can render bacterial populations virtually unlimited by mutation supply. This leads to the omnipresence and rapid generation of intraspecific trait variation, which can play a major role in ecological dynamics^14^, especially since mutations in the bacterial genome can have strong effects on traits. For instance, a single point mutation in the gene *rpsL* can make a bacterium over 250-fold more resistant to the antibiotic streptomycin^15^. Overall, strong evidence has emerged in recent decades showing that rapid evolution can alter ecological dynamics in communities across a range of systems, even those with lower rates of evolution compared to microbial systems.^16^ Similarly, a community ecology context can be important for evolutionary trajectories.^17^ Nevertheless, microbiome studies have largely overlooked evolutionary events. Intriguingly, in the case of antibiotics, while there is an extensive number of studies on both the species compositional effects of antibiotic perturbation on the microbiome and the genetics of antibiotic resistance in individual species, both aspects have rarely been analyzed together. Furthermore, high-throughput sequencing based approaches allowing the investigation of mutations in longitudinal microbiome data have focused on *in vivo* and field samples where the species are uncultured and the pre-existing traits of the species cannot be explicitly estimated^18^. This confounds the ability to assess the importance of evolution relative to species sorting.

Ecological resilience describes the ability of a community not only to withstand a perturbation (ecological resistance) but also to recover from it. Studies have shown variable results concerning the resilience of the human gut microbiota to antibiotic perturbations at high, clinical concentrations^19-21^. Effects of antibiotics that may or may not completely reverse include reduced community stability^22^ and diversity^23^, which have been associated with adverse health consequences in patients^24,25^. However, the sparsity of longitudinal studies and the variability in time points sampled since the perturbation pose challenges for comparing studies and assessing whether the communities are still in a state of recovery at the time of sampling. Moreover, the conditions determining a particular community response remain unclear. These include factors such as the level of perturbation and species immigration^26^, a key feature of microbial communities and form of bacteriotherapeutic (e.g. probiotic supplementation after antibiotic treatment)^27^. There is also a practical need to understand how the disturbance and immigration responses of bacterial communities interact, for example, for the design of minimal artificial communities to replace fecal microbiota transplantation (FMT), where gut microbial communities disturbed by antibiotic pulses are treated by a fecal “immigration” to restore a healthy microbiota^28^.

Here we used a 34-species model bacterial community to examine the role of ecological and evolutionary processes in the community response to different levels of pulse disturbance by the aminoglycoside antibiotic streptomycin in the absence or presence of species immigration. We performed a serial passage experiment, collecting amplicon data to track ecological dynamics and deep sequencing data to track evolutionary dynamics, and combined experimental data with pre-existing trait data on community members (Figure 1). We found that communities responded sensitively and repeatably to the different environments, which was linked to species sorting by pre-existing phenotypic traits relevant to fitness and increasing species extinctions as a function of antibiotic level. Adaptive mutations also occurred but could not be linked to the ecological dynamics. Despite the sensitive response to the perturbation, communities were able to recover close to the initial community state in all but the highest antibiotic level. However, the loss of species as a function of antibiotic level as well as the occurrence of evolutionary changes within species still left persistent changes in communities, compromising their resilience over the long term. Importantly, immigration played a key role in resilience at the species level by preventing species extinctions.

**Figure 1.**
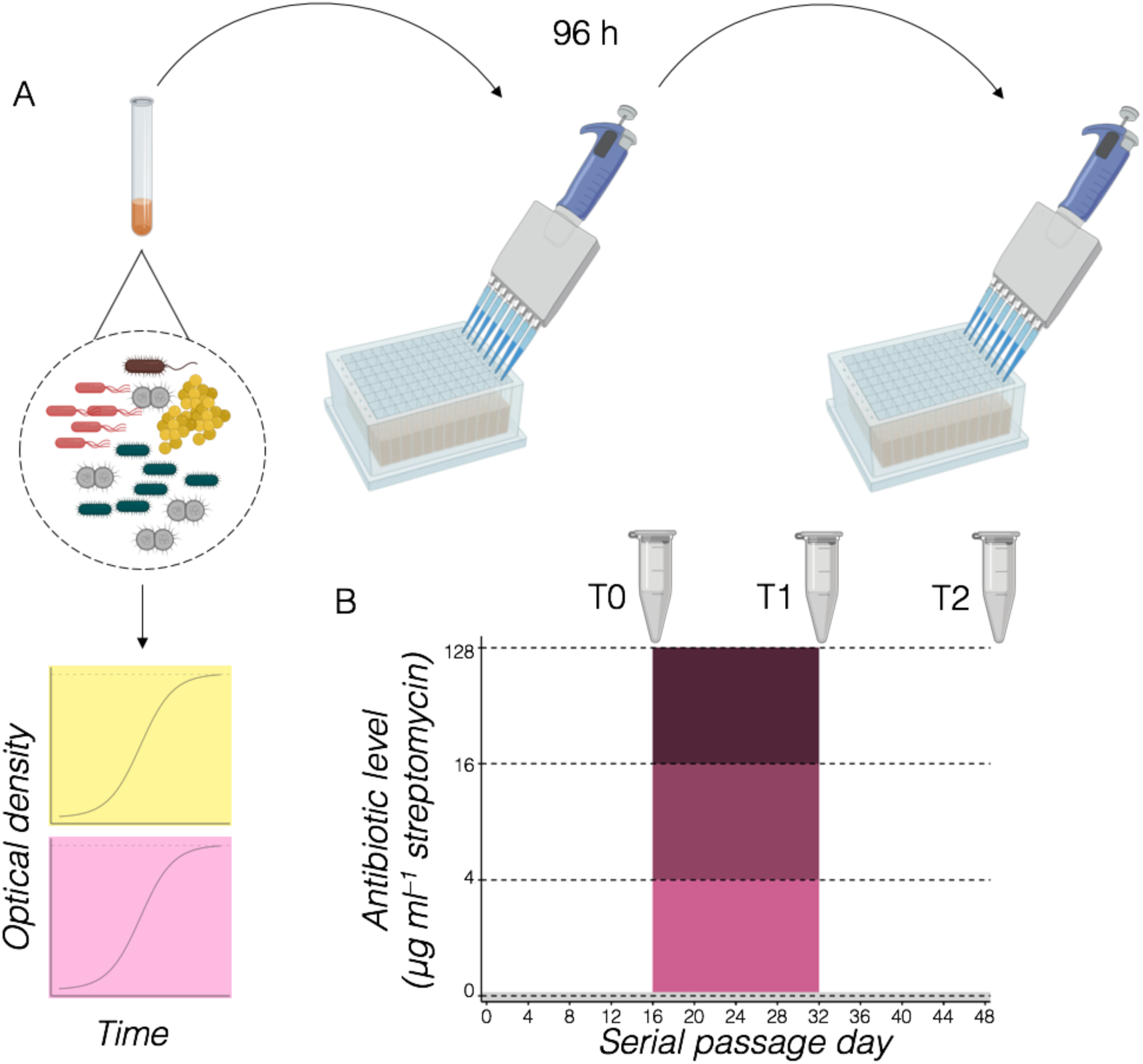
Experimental design. (A) Physical setup. A serial passage experiment was conducted with a 34-species artificial community in deep 96-well plates. Initial key traits of community members (intrinsic growth rate, represented by growth curve with yellow background, and intrinsic antibiotic susceptibility level, represented by growth curve in purple background) were measured separately for individual isolates used to construct the community. (B) Layout of antibiotic pulse experiment. The experiment consisted of three 16-day epochs: an acclimatization period, an antibiotic pulse period with three different levels of pulse antibiotic disturbance together with an antibiotic-free control treatment, and a recovery period. To investigate the role of species immigration, the full experiment was performed without and with reintroducing a small amount (1:500 cells relative to serial transfer inoculum) of the original community at each transfer. Each unique treatment combination was replicated eight times. Samples (N = 192) were collected for DNA extraction prior to the pulse, after the pulse and after recovery to track community composition (amplicon sequencing) and genomic evolution (metagenomic sequencing).

## Results

### Both antibiotic level and immigration strongly determine ecological resilience

The communities were compositionally sensitive to the different antibiotic levels and the presence of immigration (Figures S1, 2 & 3). Machine learning models could be trained to correctly predict the antibiotic level during the antibiotic pulse from species composition data (random forest, rf, model using community composition data from all treatments immediately post-perturbation to classify antibiotic level: permutation test *p* < 0.001, accuracy estimated by leave-one-out cross-validation, LOOCV, 0.88). However, the ability to distinguish between the antibiotic levels decreased after the recovery period (rf model using community composition data from all treatments immediately post-perturbation to classify antibiotic level: permutation test *p* < 0.001, LOOCV accuracy 0.53). In contrast, machine learning models could correctly classify the immigration treatment only after the recovery period (rf model using community composition data from all treatments immediately post-perturbation to classify immigration presence/absence: permutation test *p* = 0.19, LOOCV accuracy 0.53; rf model for immigration post-recovery: permutation test *p* < 0.001, LOOCV accuracy 0.69). These results suggest that the antibiotic perturbation had a compositional effect specific to the antibiotic level, this effect decreased with recovery, and the latter process was influenced by species immigration.

To more precisely investigate factors contributing to ecological resilience, we inspected the effect of the disturbance on two measures of entropy, Shannon diversity (information entropy incorporating species richness and evenness) after the disturbance and Kullback-Leibler (KL) divergence (relative entropy comparing species composition) in individual communities over time after the disturbance relative to the pre-disturbance state, adapting a recently suggested approach^29^ (Figure 3). An analysis of this data shows that diversity decreased (ANOVA for linear regression model on Shannon diversity with lowest AIC value; antibiotic *F*_3,60_ = 12.1, *p* < 0.001; the immigration treatment did not have a significant effect during the antibiotic pulse and was not included in the best model; Figure 3a) and community composition became increasingly altered as a function of antibiotic level during the pulse, and that immigration enhanced community recovery after the pulse (ANOVA for gls model on KL divergence with lowest AIC value: antibiotic level *F*_3,118_ = 19.8, *p* < 0.001; immigration *F*_1,118_ = 6.33, *p* = 0.013; recovery time *F*_1,118_ = 8.31, *p* = 0.005). Pairwise comparisons of estimated marginal means for KL divergence show significant differences (*p* < 0.02) between all of the antibiotic levels except for the control and lowest level.

**Figure 2.**
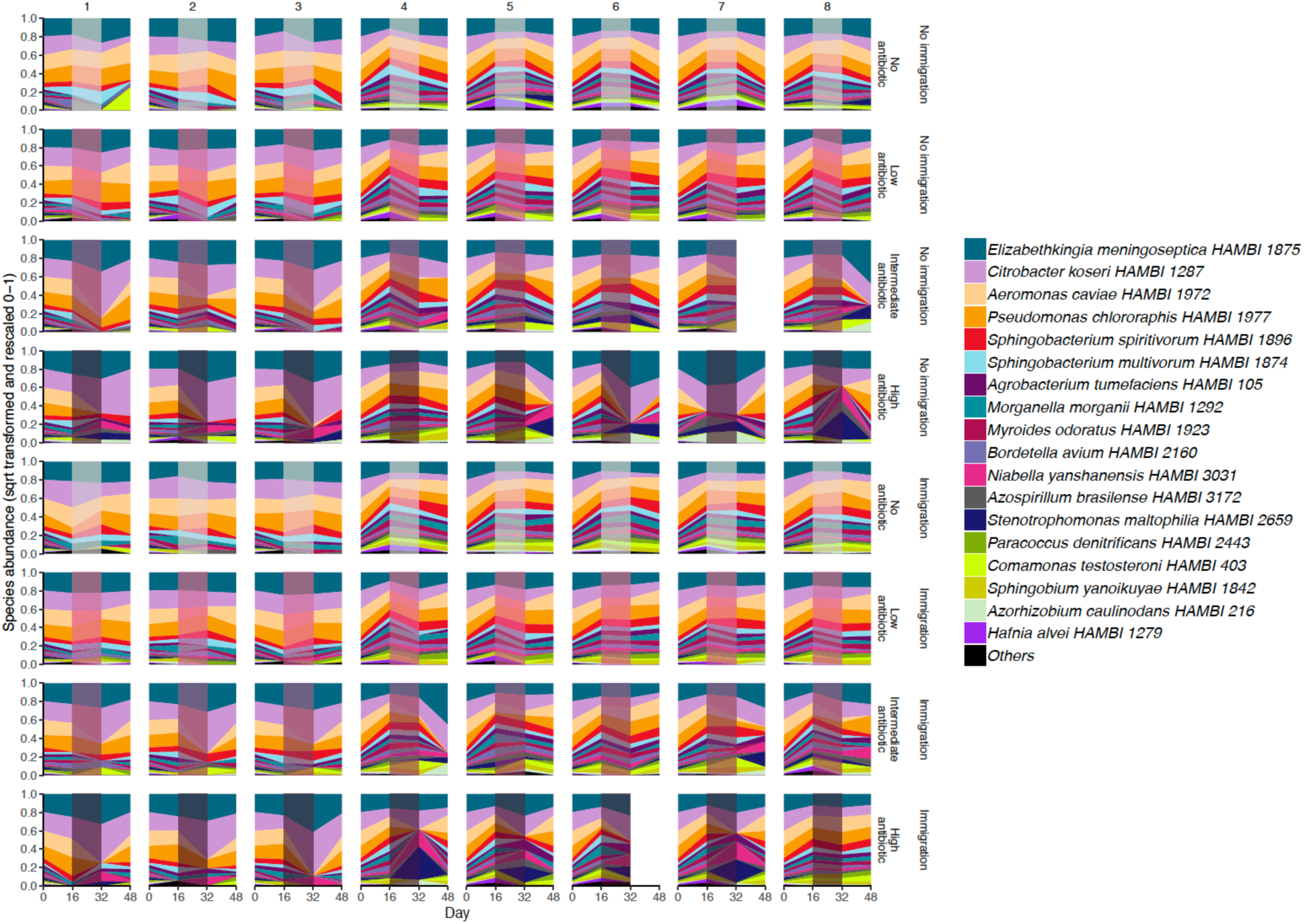
Community dynamics during antibiotic pulse experiment. The figure depicts the frequencies of abundant species across time in eight replicate communities for each unique treatment combination indicated on the right (low, intermediate and high antibiotic levels correspond to 4, 16 and 128 μg ml^−1^ streptomycin, respectively). The shaded area shows the antibiotic pulse epoch, with increasingly dark hue indicating increasing antibiotic level. The top four panels show the different antibiotic levels for the immigration-free treatment and the bottom four panels for the immigration. The *y*-axis has been square root transformed and scaled 0–1 to allow visual discernment of less abundant species. “Others” denotes rare taxa that fail to reach a frequency of 5 % in at least one community and time point. In total, 190 experimental samples are included in the figure together with one stock community sample to represent initial species composition for all communities. For two communities, adequate amplicon sequence data could not be recovered for day 48.

**Figure 3.**
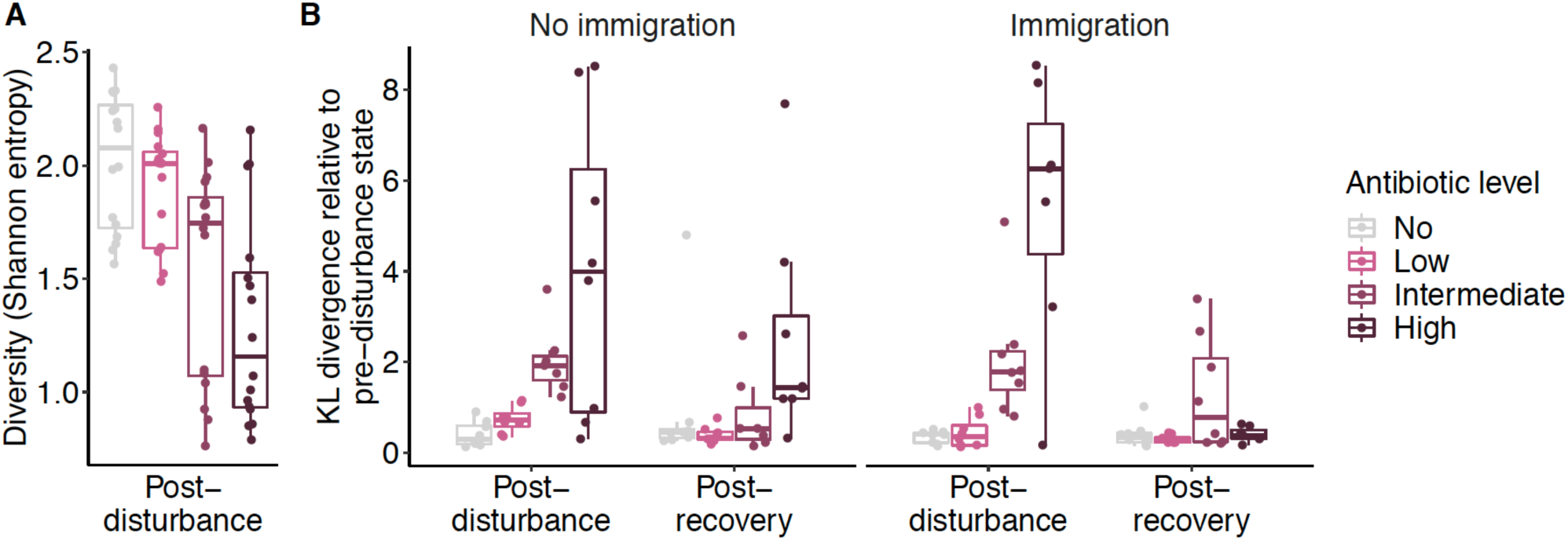
Community response to antibiotic perturbation. (A) Shannon diversity at the end of the antibiotic pulse (N = 64). (B) Ecological resilience without (left) and with (right) species immigration (N = 190). Resilience has been quantified for each community separately as the Kullback-Leibler (KL) divergence of community composition over time after the disturbance relative to the pre-disturbance state. In both panels, the data for the respective metric (Shannon diversity or KL divergence, both computed from species composition data) is displayed by a box and whiskers plot overlaid by raw data points. The lower and upper hinges of the box and whiskers plot correspond to the 25^th^ and 75^th^ percentiles, while the lower and upper and whiskers extend from the hinge to the smallest or largest value, respectively (max. 1.5 × interquartile range from hinge). Low, intermediate and high antibiotic levels correspond to 4, 16 and 128 μg ml^−1^ streptomycin, respectively.

Despite a strong persistent alteration in community composition being only observed for the highest antibiotic level in the absence of immigration (Figure 3b), increasing antibiotic levels increased species extinction probability, which was strongly counteracted by immigration (ANOVA for binomial glm model with lowest AIC value: antibiotic *χ*^2^_3,1643_ = 22.3, *p* < 0.001; immigration *χ*^2^_1,1646_ = 27.3, *p* < 0.001; species *χ*^2^_25,1618_ = 166, *p* < 0.001; antibiotic × immigration *χ*^2^_2,1615_ = 7.52, *p* = 0.057; Figure S2). Therefore, in the absence of immigration, loss of species caused persistent alterations in community composition, and the magnitude of the effect was proportional to the magnitude of the perturbation. The reported community effects of antibiotic level and immigration are unlikely to have been modulated by changes in total bacterial biomass, since similar levels of biomass were observed across the experimental conditions and time points (Figure S3). This indicates that the relative abundance of species is here a close approximation of absolute abundance, which is not always the case in microbial community studies and can have important implications for study conclusions^30^.

### Ecological dynamics are highly repeatable within antibiotic and immigration treatments

We next investigated whether increasing antibiotic levels are coupled with decreased repeatability in community trajectories, as decreased stability has been reported previously^22^ and could be a signal of alternative stable states, stochastic species extinctions or adaptive *de novo* mutations. In contrast with this expectation, we found relatively low levels of divergence in community states within each antibiotic and immigration treatment (Figure 4a). To further examine this result, we explicitly estimated the competitive fitness of each species in the community during the antibiotic pulse or recovery phase using the replicator equation from evolutionary game theory. In this approach, the frequency change of one species over time is considered to be an outcome of its fitness reduced by the average fitness of the community. We found that at increasing antibiotic levels, the competitive fitness values of specific species increased or decreased dramatically, consistent with stressors increasing fitness variance^31^, which is reflected by increasingly correlated competitive fitness landscapes (Figure 4b). During the recovery phase, in turn, the variance in competitive fitness resumes lower levels, which is seen as a reversal in the direction of the competitive fitness gradient. Therefore, the high within-treatment repeatability arises from an increasingly canalized community response in a harshening competitive environment despite increasingly dramatic compositional changes including species extinctions. We could attribute 10–13 % variation in this response during the antibiotic pulse and recovery phases to the interplay between antibiotic level and key species traits, intrinsic antibiotic susceptibility (streptomycin MIC) and intrinsic growth rate (*r*_max_; Tables S1 and S2; Figure S4). For instance, two species with high growth rate combined with high antibiotic susceptibility, *Aeromonas caviae* HAMBI 1972 (MIC = 0.75 μg ml^−1^) and *Pseudomonas chlororaphis* HAMBI 1977 (MIC = 08.0 μg ml^−1^), decreased in abundance and competitive fitness at increasing antibiotic levels (Figure 2). This, in turn, allowed the competitive release^32^ of low-abundance species with higher intrinsic resistance levels, such as *Sphingobacterium spiritivorum* HAMBI 1896 (MIC > 1024 μg ml^−1^) *Stenotrophomonas maltophilia* HAMBI 2659 (MIC = 48 μg ml^−1^). The non-linearity of these effects and important roles played by other unmeasured species traits or species interactions may explain why higher explanatory power, necessary for predictive modeling, could not be obtained for species traits despite the highly consistent ecological response.

**Figure 4.**
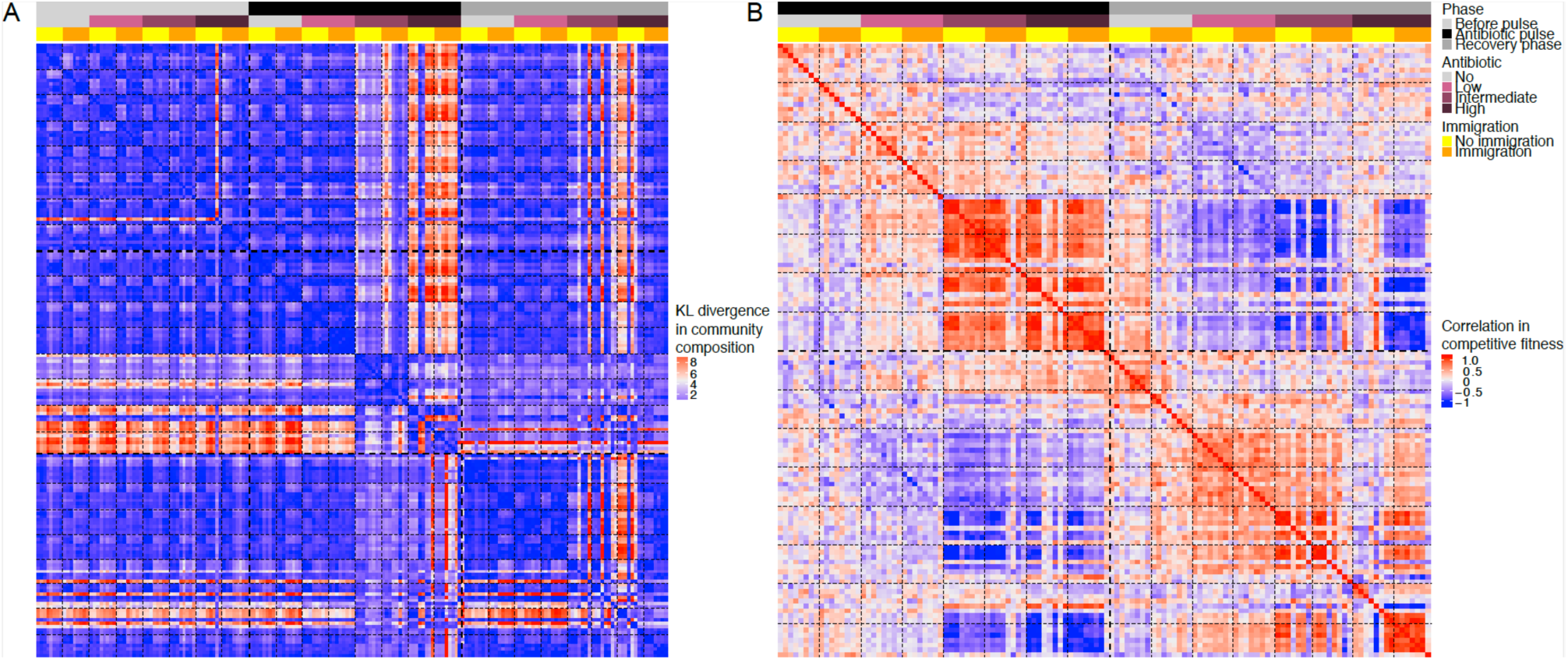
Global comparative view of community composition and competitive fitness landscape. (A) Kullback-Leibler (KL) divergence across all samples (N = 190). The color scale from blue to red indicates the degree to which community composition differs between two communities. KL divergence has been computed from species compositional data. (B) Competitive fitness landscapes across all samples during the antibiotic pulse and recovery phases (N = 126). The color scale from blue to red indicates the degree to which the competitive fitness landscapes are correlated between two communities. Correlations have been computed from competitive fitness data for each species in the communities. The heat maps have been color-annotated for the different immigration and antibiotic treatments and experimental phases. Low, intermediate and high antibiotic levels correspond to 4, 16 and 128 μg ml^−1^ streptomycin, respectively.

To more precisely inspect the low levels of divergence observed, we quantified the repeatability of community trajectories within each antibiotic and immigration treatment using the diversity dissimilarity index^33^ which relates diversity pooled over replicate communities to the mean diversity of replicate communities. This yields a value between zero and one, where zero indicates that replicate communities are identical and one that they are completely different. The community trajectories were highly repeatable (close to zero) in the different experimental treatments when species were weighted based on their abundance, although a slight (approximately 5 %) decay in repeatability was observed for the highest antibiotic level during the pulse (Shannon entropy panel in Figure 5). The high repeatability suggests that rare stochastic mutational events are unlikely to have been a major driver of the ecological dynamics of the abundant species. However, there was a higher decay in repeatability when species were given equal weight regardless of abundance, indicating that low-abundance species account for most of the loss in repeatability (species richness panel in Figure 5). When combined with the finding that the extinction probability of species increased as a function of antibiotic level, these results suggest that the stochasticity introduced by antibiotic perturbation was at least partially accounted for by ecological stochasticity caused by low-abundance species being driven extinct. Nevertheless, a potential role for evolutionary rescue in the stochasticity of the ecological dynamics of low-abundance species cannot be ruled out.

**Figure 5.**
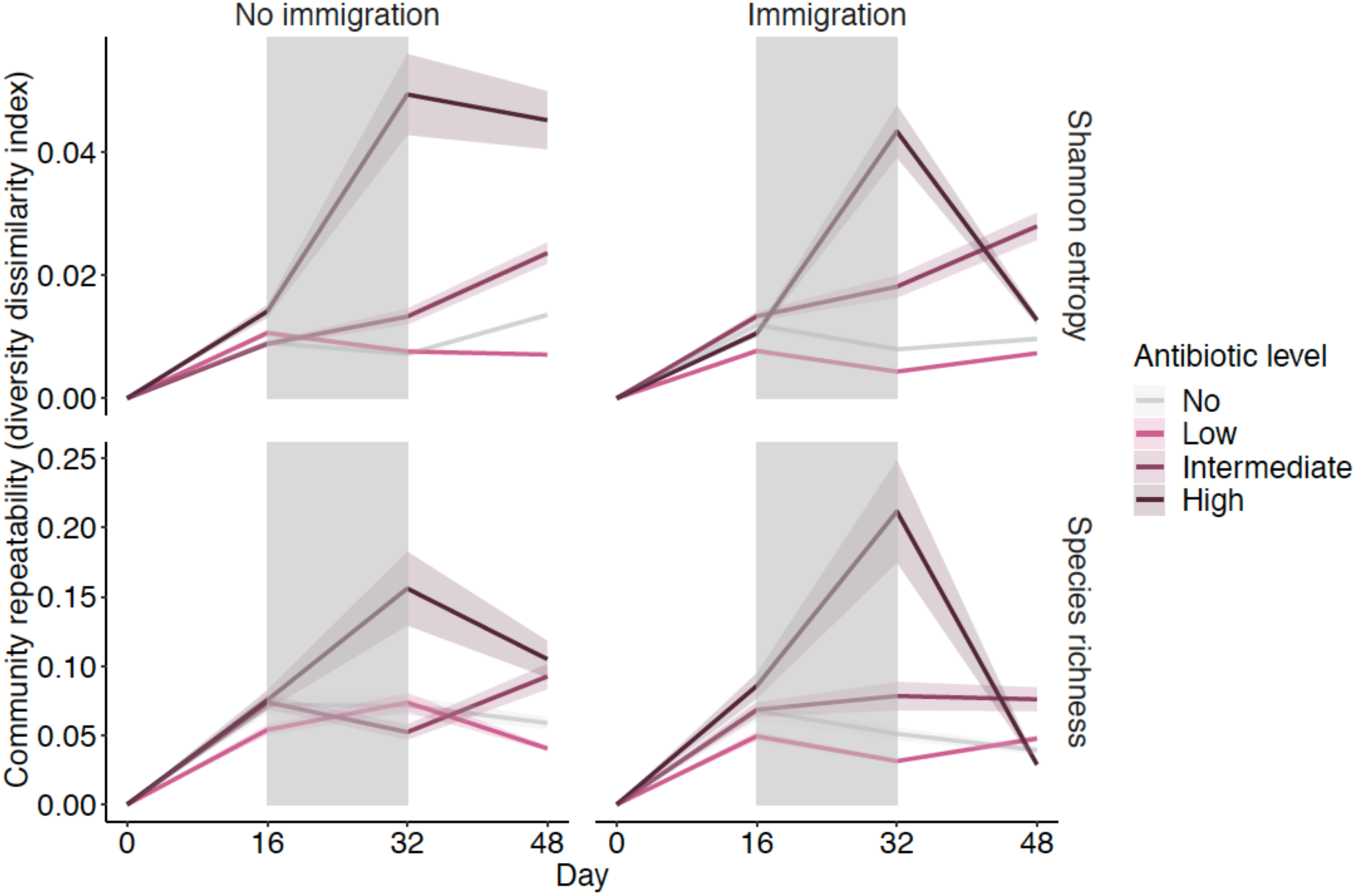
Repeatability of community trajectories, assessed using the diversity dissimilarity index (± bootstrapped standard error; N = 190). Repeatability is shown separately for Shannon diversity (top), which gives more weight to abundant species, and species richness (bottom), which gives equal weight to all species. The antibiotic pulse epoch is indicated by grey shade. Low, intermediate and high antibiotic levels correspond to 4, 16 and 128 μg ml^−1^ streptomycin, respectively. The diversity dissimilarity index has been computed from species compositional data.

### Signals of antibiotic resistance mutations occur despite repeatable ecological dynamics

To investigate mutational dynamics, we deep-sequenced three of the eight replicate communities during and after recovery from the antibiotic pulse. Most of the mutational targets estimated to be under selection (genes with more nonsynonymous hits than expected by chance) in a total of 91 genes occurred across experimental treatments, suggesting that they represent adaptations to the general experimental conditions rather than treatment-specific adaptations (Figure 6). This is incongruent with parallel mutational dynamics driving parallel ecological dynamics, since ecological dynamics diverged between experimental treatments (Figures 3 and 4). Since most of the mutational targets occurred across the experimental conditions, the mutational profiles of the communities were primarily determined by the presence or absence of whole-genome data (and thereby, information for a particular target) for the different species, although antibiotic and immigration also had minor effects on the mutational profiles (PERMANOVA for binary vectors of mutated genes: species *R*^*2*^ = 0.81, *p* < 0.001; antibiotic *R*^*2*^ = 0.015, *p* < 0.001; immigration *R*^*2*^ = 0.0082, *p* < 0.001).

**Figure 6.**
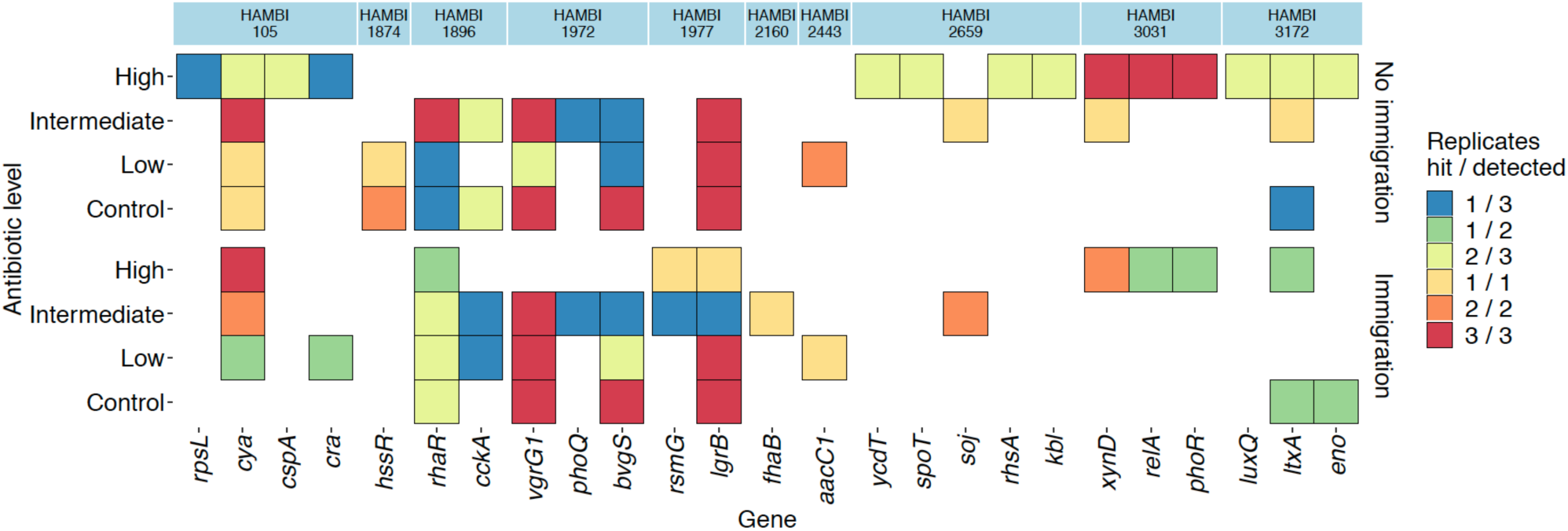
Targets of adaptive mutations reaching high frequencies (> 0.3 to fixation) in high-abundance species during or after recovery from antibiotic pulse. The same mutations were mostly observed in both time points when the species was detectable (i.e. mutation not lost during recovery). The heat map shows functionally annotated targets of recurrent nonsynonymous mutations, as well as the known streptomycin resistance gene *rpsL* which is only mutated in a single community. Since the genomic variants were recovered from deep sequencing data, they could only be confirmed for a subset of the three sequenced replicates and experimental treatments owing to differential abundance of species and volume of sequence data. Color coding is used to indicate the number of the replicates where a genomic target of interest was mutated relative to the number for which genomic variant data could be recovered in a particular experimental treatment.

Among the mutational targets that occurred only in the presence of the antibiotic, several have been previously associated with increased levels of streptomycin or aminoglycoside resistance or with the presence of these agents, suggesting that antibiotic-related adaptations occurred during the experiment. These genes include *rpsL* (encoding ribosomal protein S12, a common streptomycin resistance target)^34^, *rsmG* (ribosomal methyltransferase)^35^, *phoQ* (PhoP-PhoQ two-component regulatory system)^36^, *cya* (adenylate cyclase)^37^, *cra* (catabolite repressor/activator)^37^, *cspA* (cold shock protein)^38^, *relA* (GTP pyrophosphokinase mediating stringent response)^15^ and *spoT* ((p)ppGpp synthase/hydrolase mediating stringent response)^15^. Most of these mutations occurred in the intermediate or highest antibiotic concentration, while displaying no general pattern with respect to the presence or absence of immigration (observed only without immigration: *rpsL* (1×) and *cspA* (2×); only with immigration: *rsmG* (2×); both without and with immigration: *cra* (2×), *cya* (13×), *phoQ* (2×), *relA* (4×); Figure 6). Although the mutations occurred with antibiotic, they were not necessary for the species to be present at high abundance. For instance, an *rpsL* mutation in *Agrobacterium tumefaciens* HAMBI 105 and *rsmG* mutations in *Pseudomonas chlororaphis* HAMBI 1977 only occurred in a subset of the replicates with sufficient coverage for variant calling (indicating high abundance in the community) in the respective antibiotic treatment (Figure 6). Therefore, we could not establish a connection between the abundance of the species and the presence of adaptive mutations with the methods employed in this study. Notably, the limited number of experimental replicates deep-sequenced, limited number of species in each community with sufficient genome coverage for variant calling, and limited number of potentially adaptive mutations identified restrict the strength of inferences that can be made from the data regarding the connection between evolutionary and ecological dynamics.

## Discussion

Using a controlled setup to investigate the antibiotic response of a multispecies bacterial community in the presence and absence of species immigration, we found that the replicate communities responded repeatably to the different treatments, with the magnitude of the community response and persistent community changes increasing with increasing antibiotic levels. This was linked to increasing species extinctions at increasing antibiotic level, which prevented abundant high-growth-rate species sensitive to the antibiotic from rebounding to their pre-disturbance abundance during the recovery phase, and could be counteracted by species immigration. Furthermore, we could connect the high within-treatment repeatability of the community response to a canalized increase in fitness variance such that the noise introduced by the perturbation to the ecological dynamics was counteracted by a deterministic species response. This was explained, in part, by the intrinsic antibiotic susceptibility and growth rate of the species. Overall, these findings highlight the importance of classic ecological processes, species sorting and immigration, in defining how microbial communities respond to antibiotic perturbation.

We also found adaptive mutations sweeping to high frequencies in populations of individual species, although we could not connect these with species abundance. This shows that the disturbance response in microbial systems results from a combination of ecological and evolutionary processes operating at the same timescale. However, connecting these processes may not always be straightforward.^39^ Similar to our finding, in a recent analysis of longitudinal linked-read sequencing data from human gut microbiota subjected to antibiotic treatment, antibiotic resistance mutations were found to sweep to high frequencies in the populations of single species without necessarily resulting in an increased abundance of the species in the community^18^. In a microbial community, species sorting and adaptive mutations occur simultaneously in multiple species, and all these factors have the potential to interact, making it challenging to disentangle ecological from evolutionary processes. Furthermore, in a number of conditions, relative fitness increases within a species based on allele frequency changes do not translate into changes in absolute fitness (population size)^32^. In a multispecies setup, the competitive release of an adapted species may be suppressed, for example, by the presence of other abundant species with relatively low intrinsic antibiotic susceptibility and equal or higher resource use ability. The occurrence of a fitness trade-off between growth rate and antibiotic resistance^40^ could also make the net fitness advantage of antibiotic resistance low, causing a weak ecological effect difficult to detect in a multispecies setup. Importantly, whatever the underlying mechanism, this study supports the notion that within-species adaptive evolution can occur during a perturbation even when this is not readily suggested by the ecological dynamics.

Our findings have important implications for the understanding and management of ecological resilience. Although we found similar outcomes from antibiotic perturbation as those reported in human gut microbiome studies, such as decreased diversity^23^, these outcomes were mostly limited to the community state at the end of the perturbation, and were followed by community recovery close to the pre-disturbance state. This is non-trivial taken that priority effects^41^ or the presence of alternative stable states^42,43^ could cause a perturbed community to recover to an altered state, and emphasizes the need to assess ecological resistance during perturbations separately from recovery and longer-term resilience^44^. More generally, since the advent of amplicon and metagenomic sequencing, changes in bacterial communities have been found in response to a plethora of environmental factors, but our findings suggest that such changes may not persist and a need for caution in data interpretation in the absence of longitudinal data. Nevertheless, despite the communities mostly rebounding, species extinctions, which were more likely at increasing antibiotic levels, left persistent marks in community composition, similar to recent findings from human gut microbiota^20^, although species immigration enabled community recovery. This indicates that storing and reintroducing susceptible low-abundance species with key functionalities could play a crucial role in human management of ecological disturbances. As ecological resilience was most notably compromised for the highest antibiotic level, this suggests that in a therapeutic context, when intermediate antibiotic levels are sufficient to treat a pathogen, they may represent a desirable compromise minimizing off-target effects^3^. Notably, here we considered only a single pulse disturbance, while communities often face multiple disturbances, with historical disturbance regimes frequently priming populations, communities and ecosystems, both ecologically and evolutionarily, to similar disturbances in the future^45-47^. Therefore, the types of persistent ecological (lost species) and evolutionary (resistance mutations) changes observed in this study may have important consequences for the response of communities to future perturbations.

## Methods

### Strains and culture conditions

The liquid medium used in the experiment was specifically developed for complex communities and a long culture cycle. An artificial bacterial community consisting of 34 species (for species list, see Supplementary Materials in ^48^) was almost entirely chosen from the HAMBI Culture Collection, University of Helsinki, except for *Escherichia coli* K-12 strain JE2571^49^. The bacteria are gram-negative and represent three classes (Alpha-, Beta- and Gammaproteobacteria) in the phylum Proteobacteria and three classes (Chitinophagia, Flavobacteriia and Sphingobacteriia) in the phylum Bacteroidetes. The species are not representative of a particular natural system but were rather selected based on growth in simple, uniform laboratory conditions. Different versions of the artificial community have been used in two previous studies^48,50^, where details are reported regarding its construction and the phenotypic and genomic characteristics of the species.

A medium was specifically refined for the selected community and long culture cycles. The co-culture medium contains 1 g l^−1^ R2A broth (Labema, Helsinki, Finland) and 0.5 g l^−1^ of cereal grass medium (Ward’s Science, St Catharines, ON, Canada) in M9 salt solution. The cereal grass medium stock was prepared by autoclaving it in deionized H_2_O and filtering through 5 μl to remove particulate matter.

### Serial passage experiment

A 48-day serial passage antibiotic pulse experiment was performed consisting of three epochs: 16 days without streptomycin to allow the community composition to acclimatize to experimental conditions, 16 days with streptomycin at the concentrations 4, 16, and 128 μg ml^−1^, and 16 days without antibiotics to allow the community to recover (Figure 1). The experiment included an antibiotic-free control treatment. The experiment was performed in a full-factorial design without and with immigration consisting of adding an inoculum of the original community at each transfer. Each treatment combination was replicated eight times.

The experiment was conducted in ABgene™96 Well 2.2 ml Polypropylene Deepwell Storage Plates (Thermo Fisher Scientific, Waltham, MA, USA) in the co-culture medium. Prior to starting the experiment, all the strains were transferred to the co-culture medium and cultured for 96 hours at 28 °C / 50 rpm. Following this, they were pooled together in equal volumes and freeze-stored with 30% glycerol at –80 °C. To start the experiment, 10 μl of 100-fold diluted freezer-stock community was added to each well containing 500 μl of medium and 50 μl of sterile dH_2_O to compensate the dilution caused by streptomycin additions. The experiment was maintained every 96 hours by transferring 50 μl, about 10%, to fresh medium prepared as in the beginning of the experiment. For the immigration treatment, 10 μl of 100-fold diluted freeze-stored community was also added. For cultures containing streptomycin, the dH_2_O was replaced with an equal volume of the appropriate streptomycin stock solution.

### Data collection

The pre-existing phenotypic trait data for community members used in this study, including intrinsic growth rate and streptomycin minimum inhibitory concentration (MIC) values, was obtained as described previously^48^. To monitor bacterial density during the serial passage experiment, optical density values at 600 nm wavelength (OD_600nm_) were measured from old cultures prior to the disturbance (day 16), after the disturbance (day 32), and after the recovery period (day 48) using a well plate reader (Tecan Infinite M200 well-plate reader, Tecan Trading AG, Switzerland). Samples from time points 16 days (before streptomycin addition), 32 days (last time point with streptomycin) and 48 (final time point) days were also frozen in glycerol at –80 °C for further analysis.

DNA was extracted from the original freezer-stock community and the first three (1–3) out of eight experimental replicate communities from days 16, 32 and 48 in the serial passage experiment for a first batch of amplicon sequencing and the deep sequencing. A second batch of DNA extraction and amplicon sequencing was later performed for the remaining five replicates (4–8). DNA extraction was performed with the DNeasy 96 Blood & Tissue Kit (Qiagen, Hilden, Germany) according to the manufacturer’s instructions using 400–600 μl of sample. DNA concentrations were measured with the QubitTM 2.0 (Life Technologies Corporation, Carlsbad, CA, USA) fluorometer using the QubitTM dsDNA HS Assay Kit (Thermo Fisher Scientific, Waltham, MA, USA). Paired-end 16S rRNA amplicon sequencing (V3 and V4 regions, 2 × 300 bp; all three time points) and metagenomic deep sequencing (2 × 101 bp; only days 32 and 48) was performed by the Institute for Molecular Medicine Finland (FIMM) using the Illumina MiSeq and Illumina HiSeq2500 platforms, respectively, employing in-house protocols similar to those described before^48^.

### Sequence data processing

For 16S rRNA amplicon data, raw reads were first paired using the paired-end read merger Pear v0.9.6^51^ with defaults settings. Adapters and primers were removed from the paired reads using Cutadapt v1.10^52^ with the options -q 28 (quality-cutoff for trimming 3′end of read), -n 2 (two rounds of adapter searching), -e 0.2 (maximum error rate of 20 % for adapter identification), and --minimum-length 400 (discarding reads < 400 bp after quality control steps). Read quality was controlled with FastQC v0.11.8 (www.bioinformatics.babraham.ac.uk/projects/fastqc) and MultiQC v1.7^53^ before and after running Pear and Cutadapt. USEARCH v1.10^54^ was used to quality filter the reads using the --fastq-filter command with the options -fastq_maxee 1 (maximum expected errors 1), -fastq_truncqual 10 (truncating reads at first incidence of quality 10), -fastq_minlen 150 (minimum read length after other filtration steps), and -fastq_trunclen 150 (truncating reads at length 150 bp). Unique sequences were obtained by dereplicating using the VSEARCH v2.13.3^55^ command --derep_fulllength, followed by removal of chimeric sequences using the VSEARCH command --uchime_denovo with default settings. The reads were mapped to a reference database containing the 16S rRNA gene sequences of the 34 experimental species with USEARCH -closed_ref command with > 97 % identity requirement. Problems associated with closed reference operational taxonomic unit (OTU) clustering for environmental bacterial communities^56^, such as false positive genus assignment, should not apply to this case as the community is defined and has its own reference database. The two DNA extraction and amplicon sequencing batches (replicates 1–3 vs. replicates 4–8) display a minor but distinct batch effect in community composition (Figure 2). However, performing downstream analyses separately for the two batches did not affect the qualitative findings in the study.

For deep sequencing data, Cutadapt 1.12^52^ was used to remove sequencing adapters and quality trim sequence data, with the parameters -O 10 (minimum overlap for an adapter match), -q 28 (quality cutoff for the 3′end of each read), and --minimum-length 30 (minimum length of trimmed read). Sequence data quality before and after Cutadapt was assessed using FASTQC (www.bioinformatics.babraham.ac.uk/projects/fastqc) and multiQC^53^. The deep sequencing data was mapped to a multi-FASTA file containing the whole-genome sequences of all experimental isolates (genome accessions indicated in ^48^) except for one rare species lacking genome data (*Roseomonas gilardii* HAMBI 2470), using bowtie2^57^ with default settings. The Picard command MarkDuplicates was used to mark duplicates in alignment (BAM) files after sorting with SAMtools^58^. Subsequently, BEDtools 2.2 was used to compute genome coverage in 1 kb windows^59^. This pangenome mapping approach produced similar results compared to mapping the data to each genome individually, indicating that genome coverage was not reduced due to biased read recruitment in homologous regions, as well as cross-validating amplicon data (Figure S5). Genome coverage data also indicated a lack of major copy number aberrations (Figure S6).

Prior to variant calling and annotation, the metagenomic alignment files were split by species using SAMtools^58^. Alignment files containing below 200,000 reads, representing 5× genome coverage for a 4 Mb bacterial genome, were removed. Since differential genome coverage would affect variant count, nucleotide diversity and allele frequency estimates and thereby act as a confounder in downstream analyses comparing experimental treatments, all remaining BAM files were downsampled to 200,000 reads. Following this, genomic variants (SNPs and short INDELs) were called from BAM files with FreeBayes 1.1.0-60^60^, using a population level approach (--pooled-continuous) and calling only one variant allele per locus (--use-best-n-alleles 1). Variants were filtered based on exceeding Phred-scaled quality 20 (“QUAL > 20”) and read depth 2 (“DP > 2”) using vcffilter from vcflib (https://github.com/vcflib/vcflib). This allowed detecting variants that had reached high frequency (min. 50 %) in a total of 229 samples representing abundant species in the experiment during (day 32) and after recovery from (day 48) the antibiotic pulse. Variants were annotated using SnpEff 4.3^61^.

### Ecological analyses

All analyses were performed in the R v3.6.1 environment^62^. The t-distributed stochastic neighbor embedding (t-SNE) map for Figure S1 was created using the Rtsne package^63^ with the options perplexity = 20 and theta = 0.5. Random forest models using community composition data to classify the antibiotic or immigration treatments after the antibiotic pulse or following the recovery period were generated using the randomForest package^64^. Before analyses, rare species were removed based on > 80 % of values being zero, and the data was standardized by converting each value into a Z-score (subtracting each sample’s mean and dividing by the sample’s standard deviation). Random forest classification was performed using the function randomForest implementing the Breiman’s random forest algorithm, with the options importance = TRUE and proximities = TRUE. Subsequently, permutation tests (1,000 permutations) were implemented using the function rf.significance to test whether the models perform better than expected by chance. Following this, the function train in the package caret^65^ was used to systematically partition the data into training and tests sets repeatedly using the leave-one-out cross-validation (LOOCV) approach to estimate model performance (accuracy).

The influence of the experimental treatments on KL divergence relative to the pre-disturbance state was investigated using generalized least squares models (gls) as implemented in the nlme package^66^, specifying a residual variance structure dependent on the antibiotic level. The stepAIC function in the MASS package^67^ was subsequently used to select the best model based on the Akaike information criterion (AIC). The competitive fitness of species during the antibiotic pulse or the recovery period was estimated as the logarithm of the final frequency (rather than the change in frequency to account for the long, 16-day time interval) relative to the starting frequency, which can be directly derived from the replicator equation in evolutionary game theory. To control for noise from the frequency changes of low-abundance species and to award more weight to species with high abundance in at least one of the estimated time points, a pseudocount constituting 1 % proportion was added to the species abundance data prior to computing competitive fitness. The effect of the experimental treatments on species extinction probability was tested using the base R function glm with the option family = “binomial”.

To quantify the repeatability of ecological dynamics, we used the diversity dissimilarity index^33^:

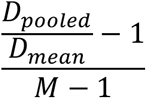

where *D* is a diversity index (either Shannon diversity or species richness computed using the vegan package^68^), and *M* is the number of communities whose species compositions are compared (over time). If the species compositions of replicates are identical, the diversity of the pooled community is equal to the mean diversity, and the diversity dissimilarity index equals 0, and if the communities have no species in common, the index equals 1.

### Evolutionary analyses

We used minimal criteria to filter raw genomic variants from the downsampled variant data prior to downstream analyses. First, we removed data for one species, HAMBI 403, which had a large number of variants (130,000) indicative of an incorrect reference genome. Second, there were peaks above 80 % in variant frequency distributions across the communities. Such a high level of parallelism suggests that the variants are either ancestral or systematic sequencing errors, and variants occurring in over 80 % of the communities were therefore removed.

From this variant data set, we extracted nonsynonymous mutations and devised a threshold for recurrence. Of all the coding genes in all the genomes, we drew mutations from a multinomial distribution with replacement. If these 588 mutations were randomly distributed over the 58,220 coding genes in the genomes, we would expect only five genes mutated in two or more populations. In total, there were 1092 coding nonsynonymous mutations across 47 genes independently mutated in two or more populations. Therefore, we focused on multi-hit genes which were independently mutated in two or more populations. This set was used for the statistical analysis below, while a larger set from genomic variant data prior to downsampling, and also including the known streptomycin resistance gene *rpsL*, was used for Figure 5 to present the maximum amount of functionally annotated potential targets of selection.

We used permutational analysis of variance (PERMANOVA)^69^ to test whether the antibiotic level or presence/absence of immigration affected the targets of mutation. Each community was scored by the presence (1) or absence (0) of a nonsynonymous mutation in each of the multi-hit genes, and these data were used to calculate the Euclidian distance between populations^70^. Before performing PERMANOVA, its assumption of homogeneity of multivariate dispersions within treatments was tested with the betadisper function in the vegan package that uses the PERMDISP2 procedure as described previously^71^. The adonis function in the vegan package was then used to test the probability that the observed distances could arise by chance by comparing them with random permutations of the raw data^72^.

### Data availability

Sequence data will be deposited in NCBI SRA. All code and pre-processed data needed to reproduce the downstream analyses and figures will be available vie Dryad/GitHub.

## Acknowledgements

We thank Aki Ronkainen and Paula Typpö for technical assistance. This work was funded by the Academy of Finland (grant 106993 to TH; grant 313270 to VM), Jenny and Antti Wihuri Foundation (grant 190040 to JC), and the Heisenberg Stipend from the German Research Foundation (DFG; grant 4135/9 to LB).

## Author contributions

Design of serial passage experiment: TH, JC, LB. Performing serial passage experiment: RJ. Design of data analysis: JC, VM, LB. Performing data analysis: JC. Interpreting results: JC, TH, LB, VM. JC wrote the first manuscript draft, with contributions from all authors. All authors approve the final version of the manuscript.

