## Supplementary Information for "Repeatable ecological dynamics govern response of experimental community to antibiotic pulse perturbation"

**This PDF file includes:**

Tables S1 to S2

Figures S1 to S5

**Table S1.** ANOVA results for effect of experimental treatments (antibiotic level and presence / absence of species immigration) and species traits (antibiotic MIC and intrinsic growth rate) on the competitive fitness of species during the antibiotic pulse period. Results are shown for the main factors and their interactions in the best model selected from the full model using stepwise model selection based on AIC.

| **Factor** | **Df** | **Sum Sq** | **Mean Sq** | ***F*** | ***P* ( > F)** |
|---|---|---|---|---|---|
| Antibiotic level | 3 | 17.5 | 5.84 | 13.3 | $1.33 \times 10^{-8}$ |
| Antibiotic MIC | 1 | 15.1 | 15.1 | 34.4 | $5.44 \times 10^{-9}$ |
| Intrinsic growth rate | 1 | 19.4 | 19.4 | 44.3 | $3.77 \times 10^{-11}$ |
| Antibiotic level × antibiotic MIC | 3 | 5.49 | 1.83 | 4.18 | $5.88 \times 10^{-3}$ |
| Antibiotic level × intrinsic growth rate | 3 | 8.59 | 2.86 | 6.54 | $2.14 \times 10^{-4}$ |
| Antibiotic MIC × intrinsic growth rate | 1 | 21.4 | 21.4 | 48.89 | $3.87 \times 10^{-12}$ |
| Antibiotic level × antibiotic MIC × intrinsic growth rate | 3 | 20.7 | 6.92 | 15.8 | $3.93 \times 10^{-10}$ |
| Residuals | 1712 | 750 | 0.438 | | |

**Table S2.** ANOVA results for effect of experimental treatments (antibiotic level and presence / absence of species immigration) and species traits (antibiotic MIC and intrinsic growth rate) on the competitive fitness of species during the period of recovery from the antibiotic pulse. Results are shown for the main factors and their interactions in the best model selected from the full model using stepwise model selection based on AIC.

| Factor | Df | Sum Sq | Mean Sq | *F* | *P* (> *F*) |
|---|---|---|---|---|---|
| Antibiotic level | 3 | 12.2 | 4.07 | 9.11 | $5.54 \times 10^{-6}$ |
| Immigration | 1 | 1.44 | 1.44 | 3.23 | 0.0725 |
| Antibiotic MIC | 1 | 12.6 | 12.6 | 28.2 | $1.27 \times 10^{-7}$ |
| Intrinsic growth rate | 1 | 13.4 | 13.4 | 30.0 | $4.97 \times 10^{-8}$ |
| Antibiotic level × immigration | 3 | 2.87 | 0.956 | 2.14 | $9.35 \times 10^{-2}$ |
| Antibiotic level × antibiotic MIC | 3 | 10.8 | 3.60 | 8.07 | $2.46 \times 10^{-5}$ |
| Immigration × antibiotic MIC | 1 | 0.159 | 0.159 | 0.355 | 0.551 |
| Antibiotic level × intrinsic growth rate | 3 | 2.48 | 0.825 | 1.85 | 0.137 |
| Immigration × intrinsic growth rate | 1 | 0.414 | 0.414 | 0.927 | 0.336 |
| Antibiotic MIC × intrinsic growth rate | 1 | 0.0271 | 0.0271 | 0.0608 | 0.805 |
| Antibiotic level × immigration: antibiotic MIC | 3 | 0.491 | 0.164 | 0.366 | 0.777 |
| Antibiotic level × immigration × intrinsic growth rate | 3 | 2.84 | 0.946 | 2.12 | 0.0962 |
| Antibiotic level × antibiotic MIC × intrinsic growth rate | 3 | 14.2 | 4.74 | 10.6 | $6.57 \times 10^{-7}$ |
| Immigration × antibiotic MIC × intrinsic growth rate | 1 | 2.02 | 2.02 | 4.52 | 0.0336 |
| Antibiotic level × immigration × antibiotic MIC × intrinsic growth rate | 3 | 2.64 | 0.880 | 1.97 | 0.116 |
| Residuals | 1642 | 734 | 0.447 | | |

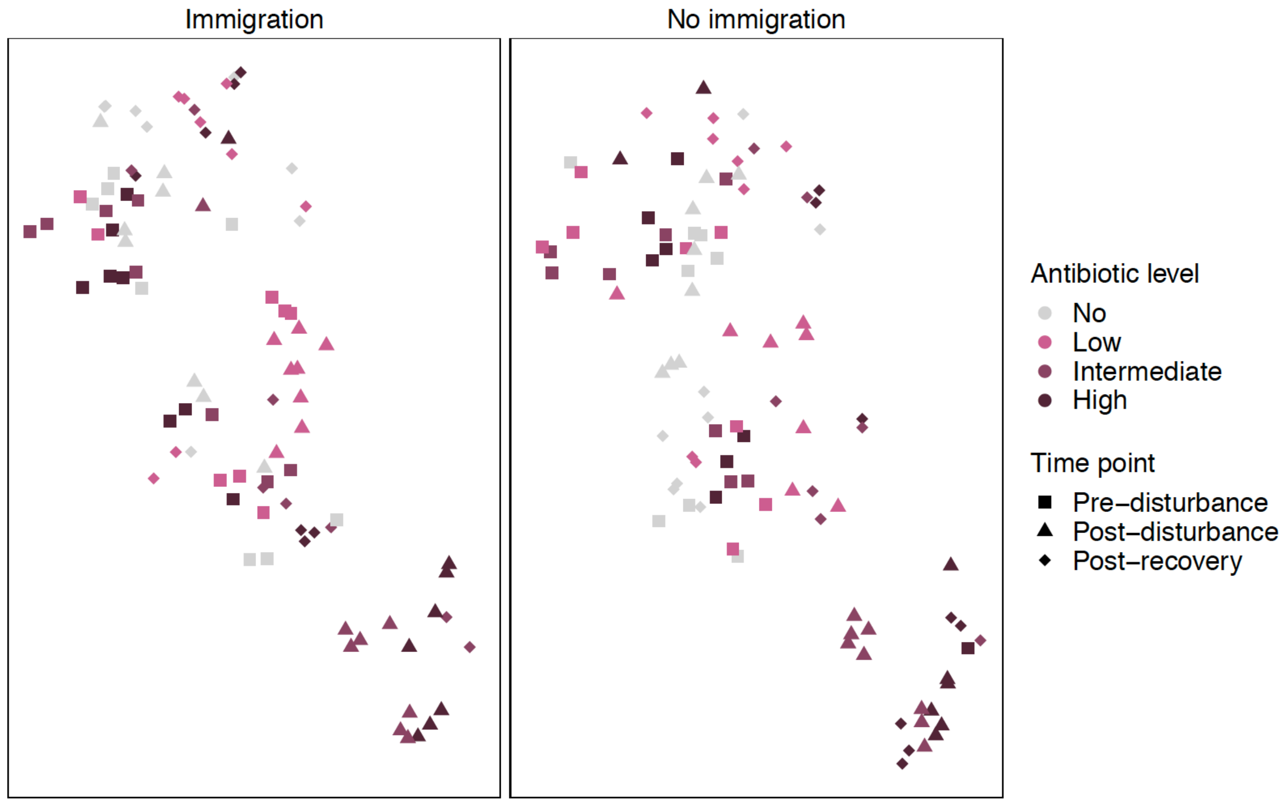


**Figure S1.** A t-SNE map showing *de novo* community clustering before, during and after recovery from antibiotic pulse at different antibiotic levels with or without immigration (N = 190). The different antibiotic levels are indicated by color coding, and the time points relative to the antibiotic pulse are indicated by different shapes. Low, intermediate and high antibiotic levels correspond to 4, 16 and 128 μg $ml^{-1}$ streptomycin, respectively. All data points originate from the same t-SNE analysis and have been separated into two panels (with same arbitrary axis units) only for the sake of visual clarity of immigration effect (at high antibiotic level, post-recovery communities indicated by triangles more often resume pre-disturbance composition in upper left-hand region).

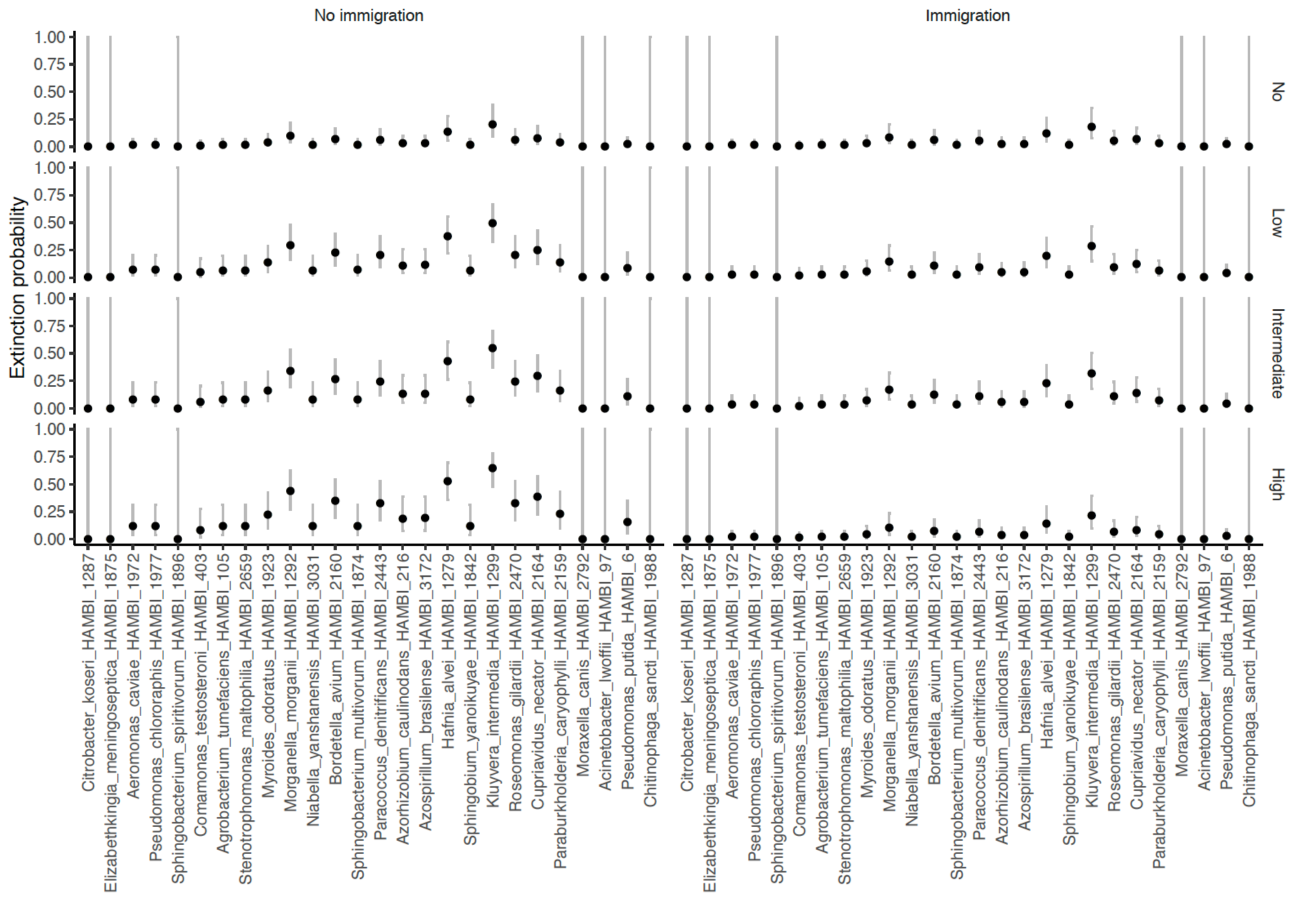


**Figure S2.** The extinction probability of species as a function of antibiotic level and the presence/absence of immigration (binomial glm estimate ± 95 % confidence intervals). Extinction is defined as the absence of a species after the antibiotic pulse (day 32 onwards) that was present prior to the pulse (day 16), and has been computed only for the species fulfilling these criteria in at least one experimental community (in total, 146 cases of extinction were observed). Low, intermediate and high antibiotic levels correspond to 4, 16 and 128 µg ml$^{-1}$ streptomycin, respectively.

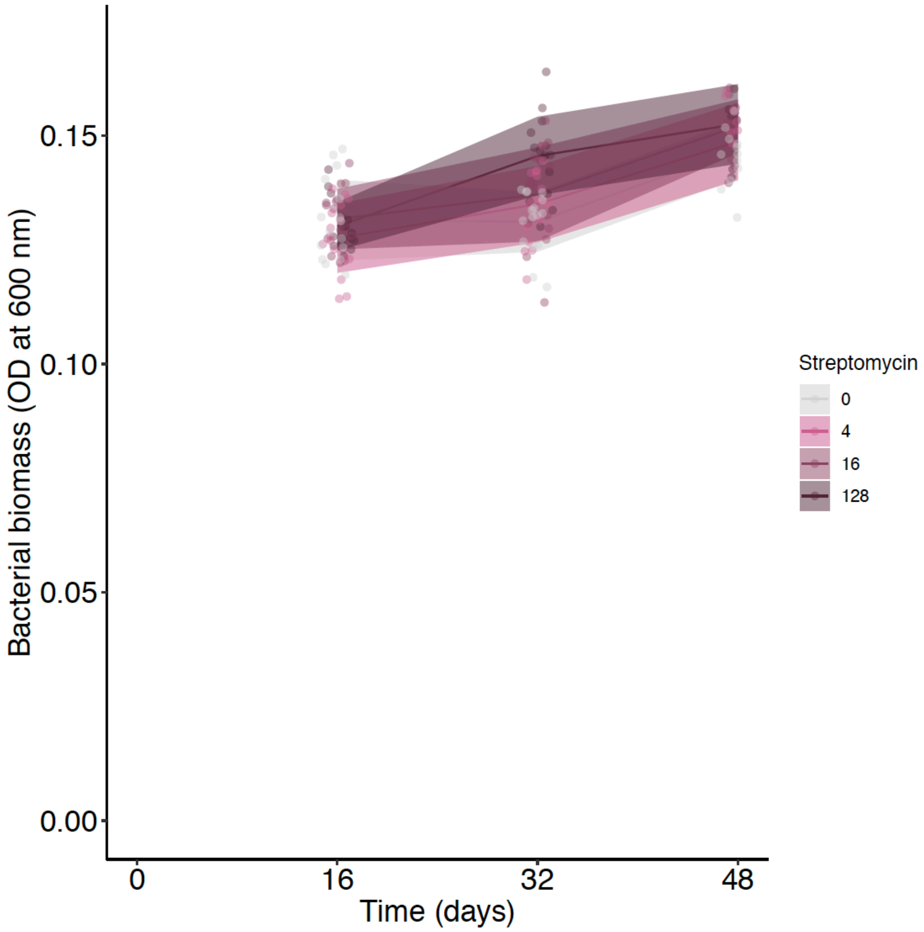


**Figure S3.** Bacterial biomass estimated by optical density (OD) at 600 nm at different levels of antibiotic pulse (expressed in µg $ml^{-1}$) in the pre-disturbance (day 16), post-disturbance (day 32) and post-recovery (day 48) phases (mean ± standard deviation).

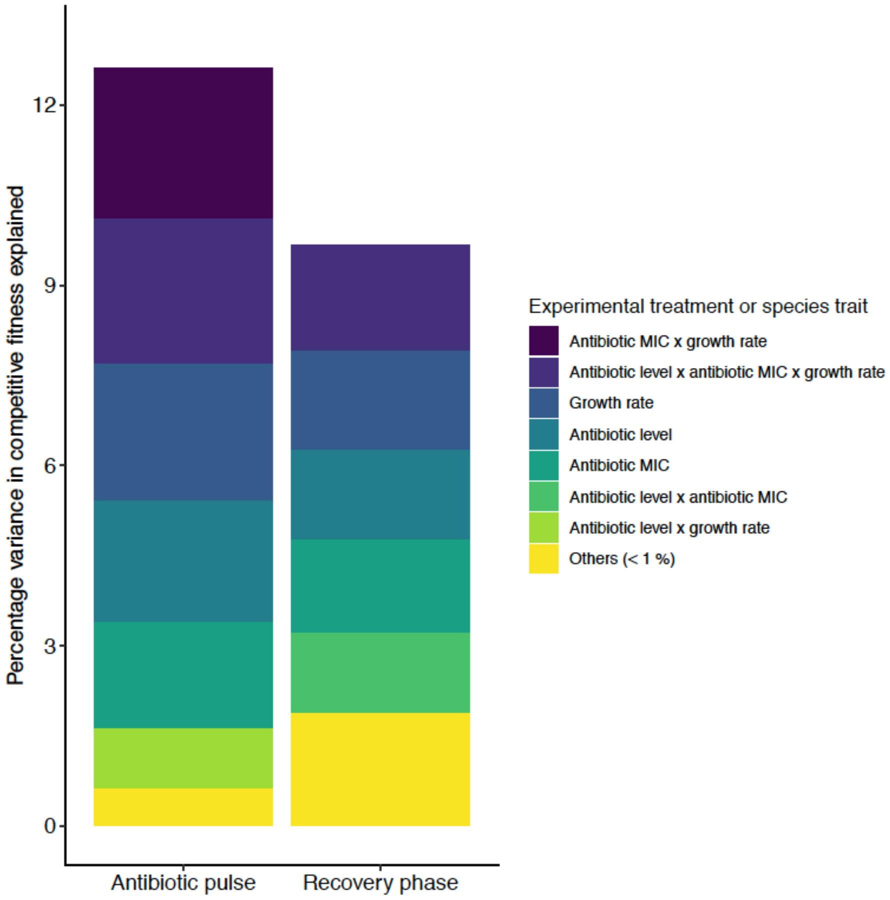


**Figure S4.** Percentage of variance in the competitive fitness of species explained by the experimental treatments (antibiotic level and presence / absence of species immigration) and species traits (antibiotic MIC and intrinsic growth rate). The variance partitioning is based on ANOVA on competitive fitness performed separately for the antibiotic pulse and recovery phases (detailed results are presented in Tables S1 and S2).

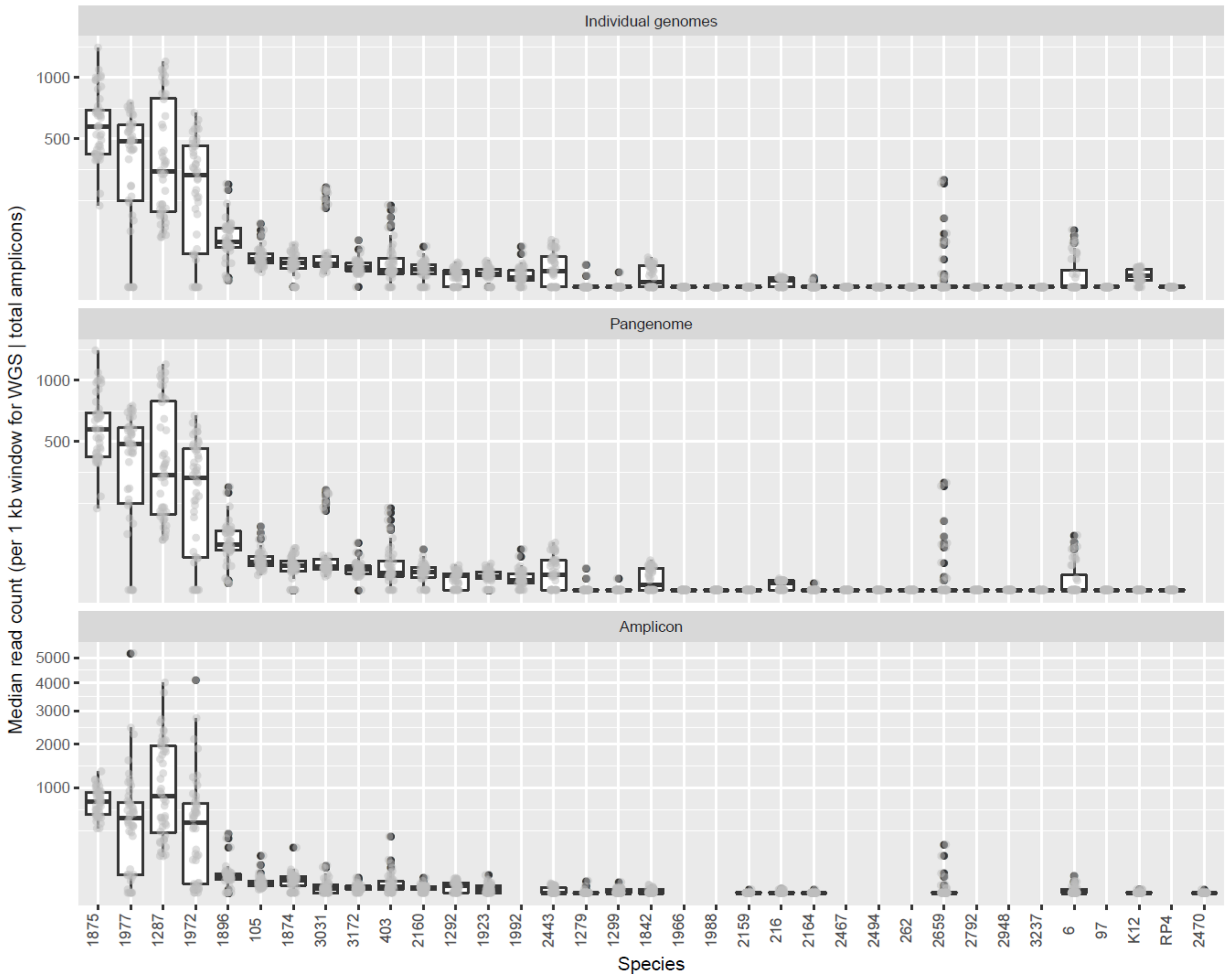


**Figure S5.** Illumina read recruitment (median per species) in whole genome alignments for deep sequencing data (two upper panels) or raw 16S rRNA amplicon data (bottom panel). The HAMBI codes of the species are indicated in the horizontal axis, with two exceptions: K12 and RP4 denote the chromosome and plasmid sequence, respectively, from *E. coli* JE2571. Read recruitment in whole genome alignments is indicated as number of reads (100 bp) in 1,000 bp blocks, and needs to be divided by 10 ($\frac{100\ bp \times read\ count}{1{,}000\ bp\ block}$) to obtain an estimate of genome coverage. For instance, a median read count of 1,000 corresponds to roughly 100× genome coverage. In the uppermost panel, deep sequencing data was mapped separately to the genome of each individual species, and in the middle panel, the data was mapped to a multi-FASTA file containing all the genomes, producing comparable results. 16S rRNA amplicon read counts have been normalized to 15,000 reads per sample.

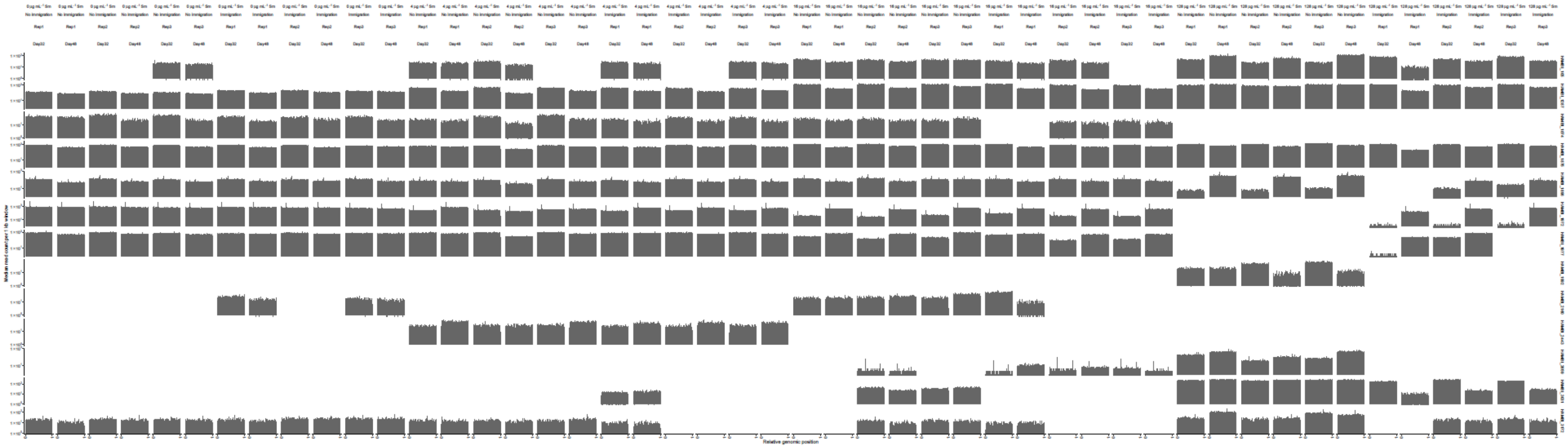

**Figure S6.** Deep sequencing read recruitment across the genomes of abundant species. Genomic position is indicated as relative position (0–1) across the whole chromosome for closed genomes or largest contig for draft genomes. Read recruitment in whole genome alignments is indicated as number of reads (100 bp) in 1,000 bp blocks, and needs to be divided by 10 ($\frac{100\ bp \times read\ count}{1{,}000\ bp\ block}$) to obtain an estimate of genome coverage.
